# CONTINUATION: Evaluation of adaptive somatic models in a gold standard whole genome somatic dataset

**DOI:** 10.1101/093534

**Authors:** Fabien Campagne

## Abstract

In http://dx.doi.org/10.1101/079087, we presented adaptive models for calling somatic mutations in high-throughput sequencing data. These models were developed by training deep neural networks with semi-simulated data. In this continuation, I evaluate how such models can predict known somatic mutations in a real dataset. To address this question, I tested the approach using samples from the International Cancer Genome Consortium (ICGC) and the previously published ground-truth mutations (GoldSet). This evaluation revealed that training models with semi-simulation does produce models that exhibit strong performance in real datasets. I found a linear relationship between the performance observed on a semi-simulated validation set and independent ground-truth in the gold set (*R*^2^ = 0.952, *P* < 2^−16^). I also found that semi-simulation can be used to pre-train models before continuing training with true labels and that this pre-training improves model performance substantially on the real dataset compared to training models only with the real dataset. The best model pre-trained with semi-simulation achieved an AUC of 0.969 [0.957-0.982] (95% confidence interval) compared to 0.911 [0.890-0.932] when training with real labels only. These data demonstrate that semi-simulation can be a very effective approach to training filtering and ranking probabilistic models.

## INTRODUCTION

This manuscript is a continuation to Torracinta et al. [2016]^1^. The reader is referred to Torracinta et al. [2016] for background and details of the adaptive deep learning concept tested in this continuation.

## RESULTS

Following our earlier presentation of deep-learning methods to train probabilistic models for somatic variation calling, I evaluated the performance of adaptive models with data from the International Cancer Genome Consortium (ICGC). The ICGC recently published a benchmark dataset: the ICGC GoldSet Alioto et al. [2015].

The ICGC GoldSet consists of data from a matched normal and tumor sample, which both have been subjected to high coverage sequencing (e.g., about 300x). The high-coverage data were used by members of the Alioto study to determine the ground truth of somatic variation in the tumor sample. Using these data, new somatic mutation calling approaches can be evaluated in the reduced coverage datasets using ground-truth variations. A drawback of the ICGC GoldSet evaluation protocol is that some mutations with low frequencies (e.g., 10%) that are visible in the 300x data can be undetectable in the reduced coverage datasets. Such mutations are labeled as “GOLD” only in Supplementary Table 1 of Alioto et al. [2015], because they were called only in the high-coverage dataset and could not be identified by any caller in the normal coverage dataset.

To evaluate semi-simulation, I trained adaptive models using the ICGC GoldSet normal and tumor samples (see Methods). In the absence of two germline samples, I used the tumor sample as the sample in which semi-simulation plants mutations. The drawback of this training protocol is that the probability of mutation can be slightly underestimated at the true mutation sites. The model was trained with a random sample of 10% of the sites sequenced by ICGC. Site sampling was random and made no attempt to include the true mutation sites described by ICGC (in supplementary material of Alioto et al. [2015]). If some true sites are included in the semi-simulated dataset, their label is completely controlled by semi-simulation, and not influenced by the GoldSet ground-truth. Table 1 provides a description of the training, validation and test sets used to train semi-simulation models with these data (ICGC-10 semi).

**Table 1.** Dataset Characteristics

|  | # Training Examples | # Validation Examples | # Test examples | # Features |
| --- | --- | --- | --- | --- |
| ICGC-10 semi | 21,137,888 | 1,172,049 | 1,173,442 | 280 |
| ICGC gold-set | 37,920 | 18,690 | N/A | 280 |

The model trained on ICGC-10 obtained an AUC of 0.9581 on the validation set and a test set AUC of 0.955 (95% confidence interval [0.951-0.959], calculated using 10,000 random examples from the test set, see Methods).

To determine if such a semi-simulation trained model can be predictive on a real dataset, I evaluated the performance of the model on the ICGC gold-set dataset (labeled ICGC gold-set in Table 1). The model obtained an Area Under the ROC Curve (AUC) of 0.883 [0.870-0.896] 95% confidence interval. While the model suffers a drop in performance on the real dataset, it is clearly predictive despite having been trained only with simulated labels for this dataset. Figure 1 shows the Received Operating Curve (ROC) for this model on the GoldSet (left) and the reliability diagram (right). The ROC indicates that the model performs well for a majority of true mutations in the ICGC GoldSet. The reliability diagram shows that predictions assigned a strong probability by the model (e.g., > 0.90) have an 85% chance of being true positives. These performance measures are similar, albeit not directly comparable, with the ones reported for state of the art somatic mutation callers in Alioto et al. [2015].

**Figure 1.**
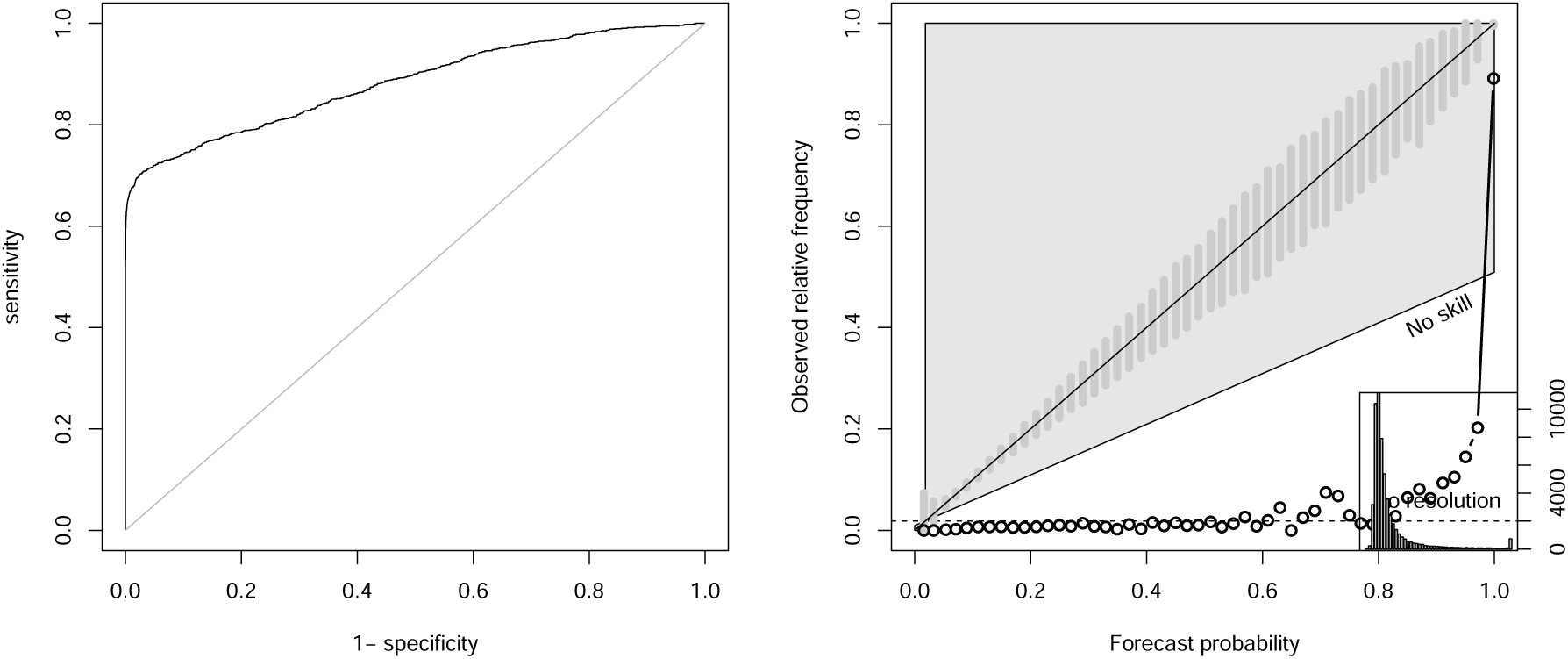
Receiver Operating Characteristic (ROC) Curve and Reliability Diagram for model predictions on GoldSet. A model trained exclusively by semi-simulation (simulated labels only) is evaluated on gold standard ground truth from the ICGC GoldSet (Alioto et al. [2015]). Left presents the ROC curve. Right presents the reliability diagram. Forecast probability is the probability generated by the model. Observed relative frequency is the proportion of true labels in a set of sites. Both plots indicate that the model performs extremely well for a majority of sites (corresponding to about 65% sensitivity), then has degraded performance and fails to identify some true positive sites described in the ICGC GoldSet. Despite the drop in performance such a model is suitable for prediction in a real dataset because strong performance is obtained for sites with highest forecast probabilities.

To better characterize how performance of semi-simulated models translate to the GoldSet, I generated a number of alternative models with random hyper-parameter choices. As usual when sampling hyper-parameters, a full range of performance is expected, from non-predictive models (AUC close to 0.5) all the way to close to the performance of the best model that can be derived from the dataset, but including models of intermediate performance. Figure 2 presents the performance of these alternative models on the GoldSet. This figure shows an almost linear relationship between performance estimates obtained on the semi-simulated ICGC-10 validation set and performance on the GoldSet (for models which were trained exclusively on ICGC-10 with semi-simulation). These data confirm that semi-simulation can help train models that perform well on a similarly distributed real dataset. Furthermore, the plot establishes that validation performance on the semi-simulated dataset can be used as a guide for selecting a model expected to perform well on a real dataset.

**Figure 2.**
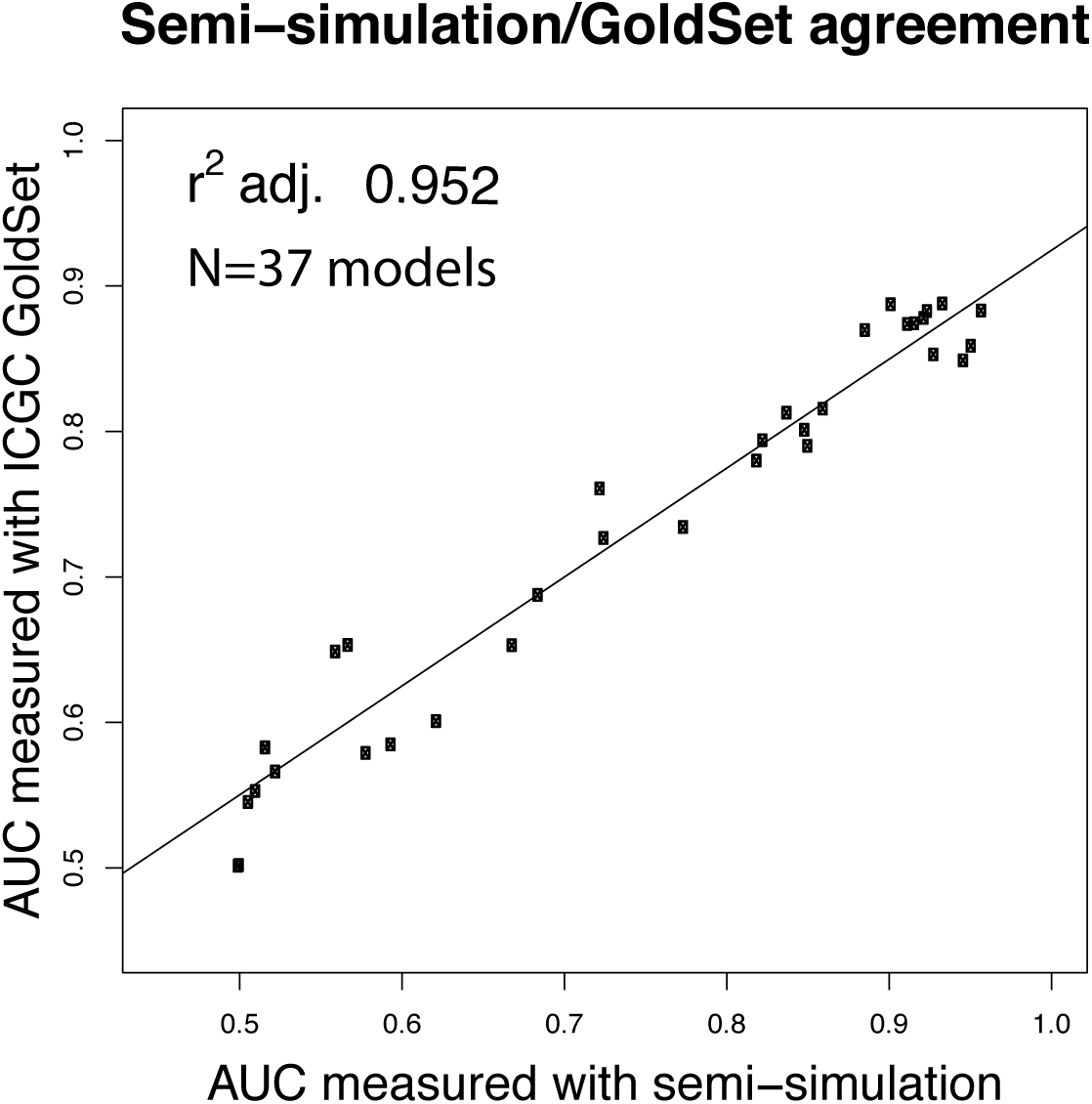
Performance of alternative models. Alternative models can be constructed by varying hyper-parameters of the training procedure (number of training examples used, learning rate, dropout rate, L2 regularization rate). Most of the alternative models are expected to have sub-optimal performance. This plot compares the performance of alternative models obtained on the validation set to the performance obtained on the GoldSet. The strong linear fit (*R*^2^=0.952, *P* <2^−16^, N=37 alternative models) with a slope of 0.75 indicates that hyper-parameter search on a semi-simulated dataset can guide model selection even in the absence of a real dataset.

Semi-simulation does not require true labels, and none were used for training the models presented so far. However, an interesting question is whether performance of semi-simulated models can be improved in cases where some amount of real labels is available. In this case, a fully trained semi-simulated model can continue training with a dataset containing real labels. In practice, this can be accomplished by loading the parameters of a trained model and resuming training with a new training and validation set (with labels from a real dataset). I have tested this scenario in Figure 3. It compares the performance of about 50 alternative models (sample of hyper-parameter choices) when the model is trained exclusively with the labeled data (Direct training with GoldSet, left), or when the model is trained with the semi-simulated data (ICGC-10, Pre-training), then re-trained with real data (GoldSet, right). Figure 3 shows that pre-training with semi-simulated data helps find many more alternative models with strong performance than is possible when training directly with labeled data from the GoldSet. This can be explained in large part because the models have about 1.6 million parameters, and are challenging to train in a dataset with only about 38,000 examples, of which only 700 are identified as somatic variations in the training set. In this case, pre-training with semi-simulation likely helps optimize most parameters of the model that do not need to be adjusted when the second training set is presented. As a result, the best models are obtained with pre-training with semi-simulation (e.g., 0.969 [0.957-0.982]) compared to 0.911 [0.890-0.932] with direct training.

**Figure 3.**
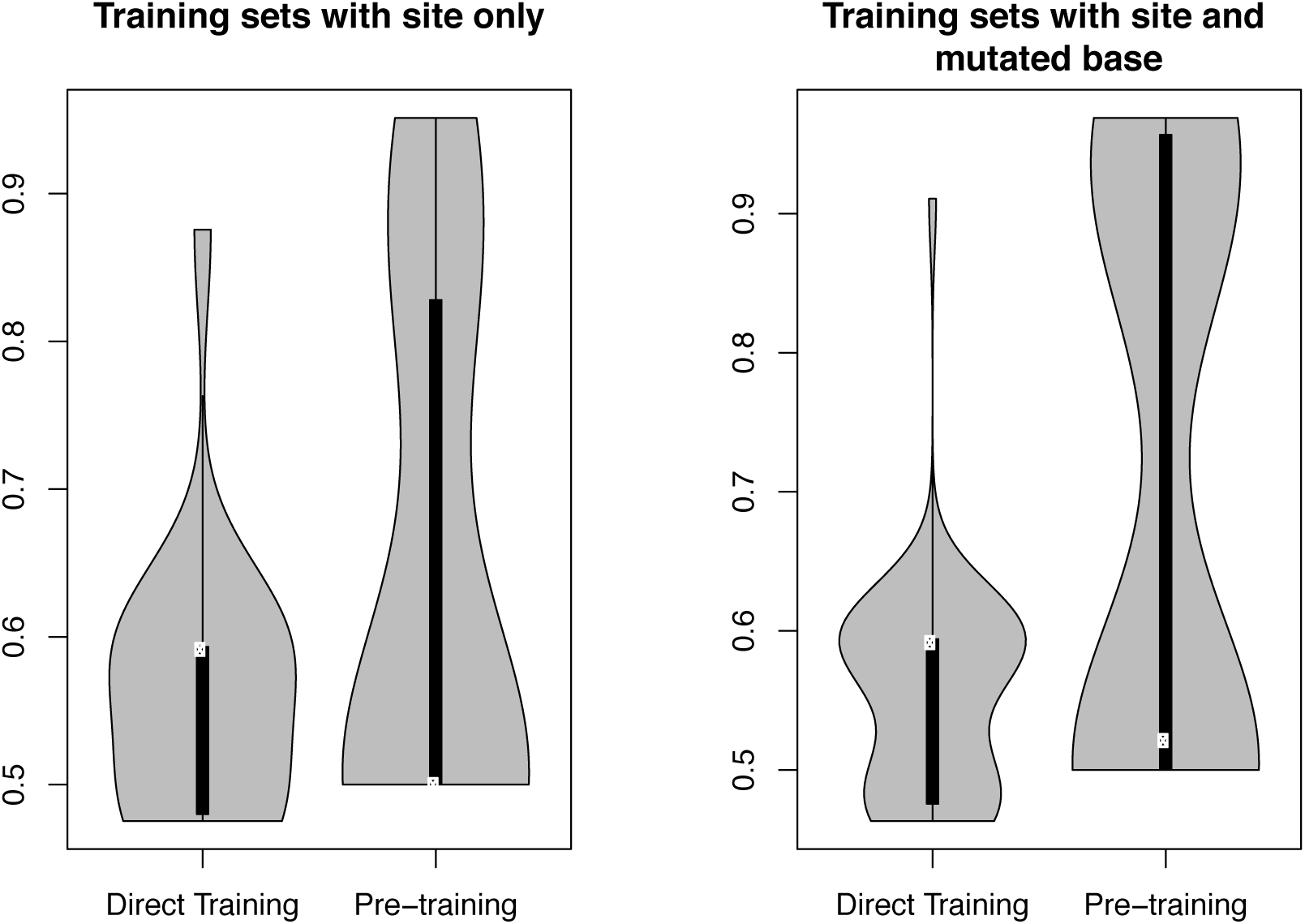
Impact of semi-simulation pre-training on labeled training. This experiment looks at the impact of pre-training with semi-simulated datasets on the validation performance of trained models. Each panel shows two violin plots of the model validation AUC. Left: direct training with labeled data from GoldSet. Right, models pre-trained with semi-simulated data before additional training with GoldSet. All models trained to initial convergence (first decrease in validation AUC stops training). I repeat the comparison for two kinds of datasets, one where only the location of the site is used to indicate that the site is a somatic mutation, the other where the site and identity of the base mutated are used to train the model. In each experiment I find that pre-training greatly increases the proportion of high-performing models. The standard error on each AUC estimate for high-performance models is not shown, but is reported in the text for the top performing models. White dots indicate the position of the larger density in each violin plot. Width of the grey shapes indicates the density of points in this region of AUC values.

## DISCUSSION

While training models with semi-simulated data may appear to train models with no supervised data, and learn something from nothing, this is not accurate. Semi-simulation relies on an understanding of the process that generates the labels, to simulate true signal and plant it in real, noisy, datasets. Semi-simulation therefore substitutes a conceptual model of a process, implemented in a simulation tool, to the usual observations of labels used so far for supervised learning. Semi-simulation is expected to help in cases where the process that generates the signal is sufficiently well understood that reasonably realistic simulations can be developed. Simulation of somatic mutations is one such problem where simulating mutations is orders of magnitude more cost effective than developing benchmark datasets to identify true mutations. Semi-simulation is therefore expected to be useful in applications where conceptual models are developed (e.g., in scientific research). It would be much less useful in applications of deep learning to domains where collecting labeled data is more cost-effective (e.g., face recognition in pictures, reinforcement learning to learn to play games).

Taken together, these results indicate that models trained with semi-simulation can yield competitive ranking and filtering approaches for genomic datasets. This evidence is important because semi-simulation makes it possible to develop models for specific assays and analysis protocols, which can adapt to the noise characteristics of assay and analysis methods, as we have illustrated in our first report about semi-simulation Torracinta et al. [2016].

I also showed that pre-training models with semi-simulated datasets can help train more predictive models. This result is important because it suggests that semi-simulation can be used not only to train models when large amounts of labeled data are not available, i.e., such as for a new assay, but also when labeled data starts to become available. We can therefore envision developing models for new assays with semi-simulation only, and iteratively refining the models as more labeled data are produced (e.g., by independent experimental validation of results ranked by prior iterations of the models). I anticipate that this iterative model development approach will yield state of the art filtering and ranking models for many assays.

## METHODS

### GoldSet sample processing

ICGC GoldSet samples for normal and tumor samples were downloaded from the European Genomics Archive (EGA) using accession code EGAD00001001859 in the FASTQ format.

Reads were converted to the Goby compact-reads format Campagne et al. [2013]: EGAD00001001859- LA-tumor 619,412,062 reads, 96.5 GB, EGAD00001001859-LA-normal 456,984,733 reads, 72.6 GB.

Compact-reads were uploaded to an internal instance of GobyWeb Dorff et al. [2013]. Reads were aligned with GobyWeb using bwa-mem, implemented in the BWA _MEM _ARTIFACT Gob-yWeb plugin (https://github.com/CampagneLaboratory/gobyweb2-plugins/tree/plugins-SDK/plugins/aligners/BWA_MEM_ARTIFACT). This process produced two alignments in the Goby format Campagne et al. [2013].

### Semi-simulation

Normal and tumor alignments were processed with GobyWeb Dorff et al. [2013]. We used the Goby Web Sequence Base Information plugin (**SBI**, https://github.com/CampagneLaboratory/gobyweb2-plugins/tree/plugins-SDK/plugins/analyses/SEQUENCE_BASE_INFORMATION) to produce raw and semi-simulated mutated .sbi files. The SBI plugin uses version 1.1 of the variationanalysis project (version 1.0 was described previously Torracinta et al. [2016]). The plugin was configured to realign reads around indels, call indels and keep sites with at least one base supporting a variation and keep sites with a single distinct read index.

The OneSampleCanonicalSimulationStrategy was used for semi-simulation, which considers sites canonical when the germline site has up to two alleles with more than 90% of bases. The plugin was configured to randomly sample 10% of sites across the genome to yield a semi-simulated training set with 23,483,379 genomic sites. This set was randomly split into a training set with 21,137,888 sites, a validation set with 1,172,049 and a test set with 1,173,442 sites. The normal sample was marked as germline and the tumor sample was marked as somatic (where mutations will be planted by semi-simulation).

### Feature Mapper

The feature mapper used in this work extends that presented in Torracinta et al. [2016] and maps information about the genomic context of a site (10 bases before and after are one-hot encoded), the density of insert sizes, read quality and mapping qualities at the site. We used implementation org.campagnelab.dl.somatic.mappers.FeatureMapperV25 (see Campagne and Torracinta [2016])

### Training with Semi-Simulation

Learning rate was set to 5 and training performed with the Adagrad optimizer, which decreases learning rate for each parameter independently during training. Other hyper-parameters for the model were searched with the search-hyper-params.sh tool in the variationaanalysis project (release 1.1+) to determine the dropout rate and regularization rate that maximizes AUC on the first 10,000 sites of the validation set (training models with the first 10,000 sites of the training set). The same model architecture as presented in Torracinta et al. [2016] was used for training models in this continuation. Training with the full training set was performed on an NVIDIA GPU GTX 1080 with early stopping, using this command line:

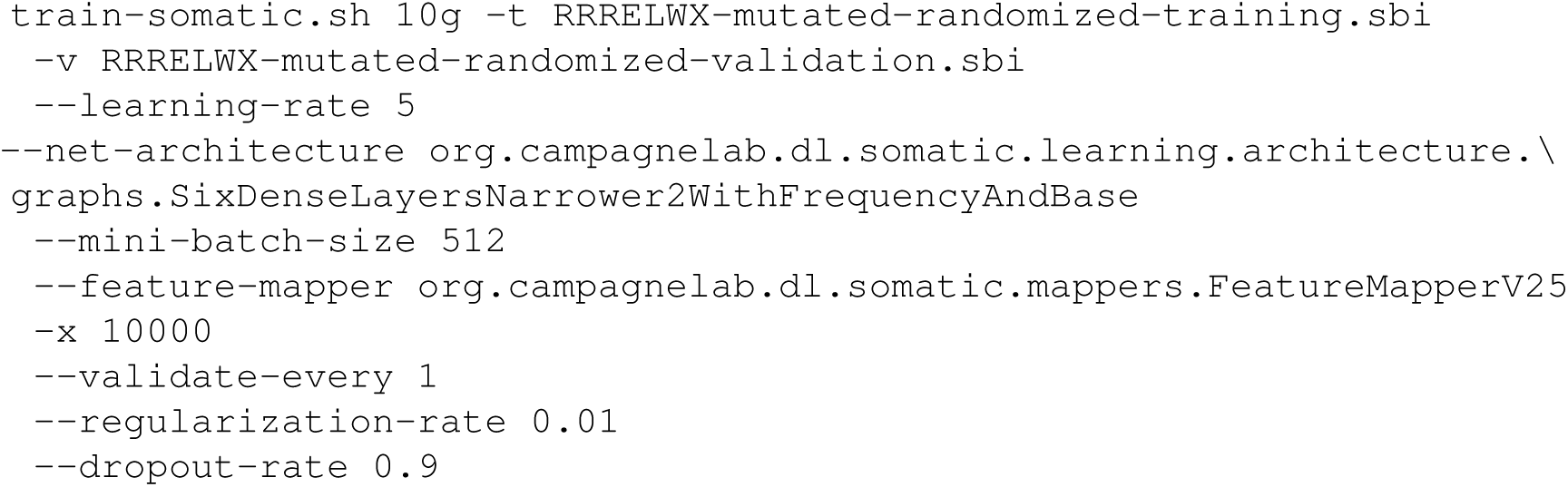

### Hyper-parameters searches

Models with various hyper-parameters were produced with the search-hyper-params.sh tool provided in version 1.1.1 of the variationanalysis project (Campagne and Torracinta [2016]). For instance, for Figure 2, the following commands generated 100 models with different learning rate:

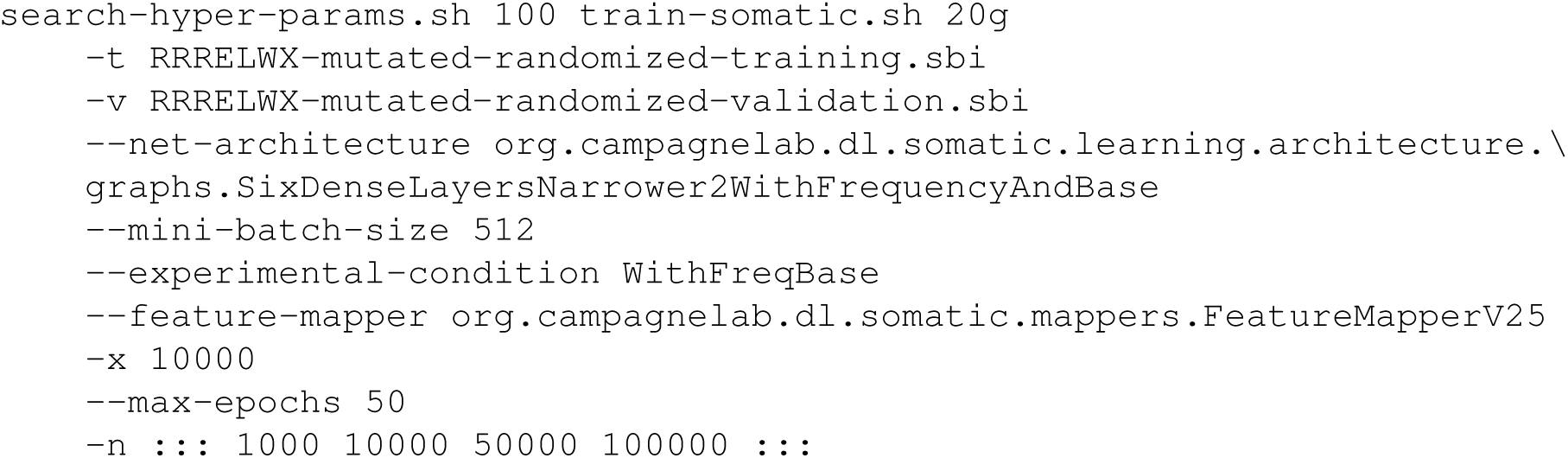

### Gold Standard Data Set

A Gold Standard dataset was constructed with the GobyWeb SBI plugin and the GoldSet annotations (SNV and indels) to yield a dataset with 79,637 genomic sites of which about 1405 sites are annotated as mutated and are true mutations in the ICGC gold set (supplementary table to Alioto et al. [2015]).

### AUC estimations

AUC, or Area Under the Receiver Operating Curve (ROC), was estimated by the exact method, by calculating the number of pairs of positive and negative examples where the positive example scores higher than the negative, and dividing by the number of pairs. Standard error of the AUC was estimated with the method of Hanley and McNeil [1982]. 95% Confidence intervals were derived by adding and substracting 2.96 times the standard error to the AUC estimate. This calculation is implemented by the variationanalysis predict.sh tool. When evaluating performance for several models, I used the predict-all.sh tool of variation analysis. For instance, to estimate AUC of models shown in Figure 2, the following command was used:

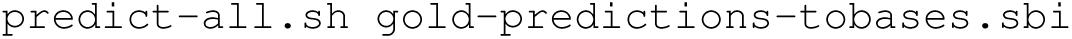

The previous command scans the models defined in the model-conditions.txt file and evaluates the performance of each model again the dataset provided in argument.

## Footnotes

1 A continuation is a preprint that continues where an earlier preprint left off. The term can also be used to refer to the initial preprint and one or more continuations of the preprint. The title of a continuation starts with the DOI of the first preprint in a continuation, followed by the word CONTINUATION in uppercase and a colon. A short sentence summarizes the results presented in the continuation. Authors of a continuation should be listed who have contributed to the material presented in the continuation, rather than to the original preprint (since these authors received credit in the first preprint already). Instead of repeating introduction and methods that are common with the prior preprint, or revising the initial preprint and force readers to read old material to discover new one, this format encourages brevity of reporting. New results or changes to methods are reported in a continuation. An important advantage of the continuation format is that it makes it possible to report results chronologically in preprints, and clearly expose the steps taken during a research study. A manuscript submitted for publication may later show only a subset of the results presented in these preprints, and may change the order of results in its presentation, in order to improve clarity for readers who encounter the ideas for the first time. Since the article can cite the preprints, it is understood that chronology is described accurately in the continuation format, while the peer-reviewed article is a simplifying summary designed to distill the key elements of a new scientific contribution.

